# Optimizing Dosage-Specific Treatments in a Multi-Scale Model of a Tumor Growth

**DOI:** 10.1101/2021.12.17.473136

**Authors:** Miguel Ponce-de-Leon, Arnau Montagud, Charilaos Akasiadis, Janina Schreiber, Thaleia Ntiniakou, Alfonso Valencia

## Abstract

The emergence of cell resistance in cancer treatment is a complex phenomenon that emerges from the interplay of processes that occur at different scales. For instance, molecular mechanisms and population-level dynamics such as competition and cell-cell variability have been described as playing a key role in the emergence and evolution of cell resistances. Multi-scale models are a useful tool to study biology at a very different time and spatial scales, as they can integrate different processes that take place at the molecular, cellular and intercellular levels. In the present work, we use an extended hybrid multi-scale model of 3T3 fibroblast spheroid to perform a deep exploration of the parameter space of effective treatment strategies based on TNF pulses. To explore the parameter space of effective treatments in different scenarios and conditions, we have developed an HPC-optimized model exploration workflow based on EMEWS. We first studied the effect of the cells spatial distribution in the values of the treatment parameters by optimizing the supply strategies in 2D monolayers and 3D spheroids of different sizes. We later study the robustness of the effective treatments when heterogeneous populations of cells are considered. We found that our model exploration workflow can find effective treatments in all the studied conditions. Our results show that cells’ spatial geometry, as well as, population variability should be considered when optimizing treatment strategies in order to find robust parameter sets.

## Introduction

Optimizing drug treatment and efficiently screening the effect of drugs is key to improving clinical treatments and ultimately extending patients’ life expectancy [18]. The emergence of resistant cells though is a complex phenomenon, mainly due to the complexity of biological systems [32], the interplay of processes that occur at different scales and an environment with an active role in this resistance [20, 13]. Molecular mechanisms and population-level dynamics such as competition and cell-cell variability have been described as playing a key role in the emergence and evolution of cell resistances [19]. For instance, high gene expression variability has been linked to aggressiveness in Chronic Lymphocytic Leukemia [6]. Genetic heterogeneity as well phenotype variability have both shown to be related to the emergence of cell resistance [4, 24, 32]. Furthermore, the environment has been described to have an effect on the cells’ response to drugs: 2D-cultured cell line screens failed in clinical studies [16] as cell cultures do rarely recapitulate the heterogeneity and drug sensitivity of the original tumor [17].

Multi-scale models (MSM) are a useful tool to study biology at very different time and spatial scales, as they can integrate different processes that take place at the molecular, cellular and intercellular levels [25, 26]. In the domain o cancer biology, MSMs have been used to connect cellular mechanisms underlying cancer drug resistance to population-level patient survival [34], study the role of physiologic resistance due to diffusion gradients of different nutrients and drugs [11], to quantitatively characterize pressure for invasion [3], among many other applications [25]. In general, multi-scale models provide a genotype-to-phenotype simulation framework, which is ideal for the study of *in-silico* drug screenings [9], the optimization of treatment regimes [1] and the exploration of genetic or environmental perturbations [22].

Multi-scale simulation can be used to conduct *in-silico* experiments and to generate new experimentally testable hypotheses, accelerating the discovery of new potential treatment strategies [2] Nevertheless, due to the hybrid approaches used to describe multi-scale models (e.g. discrete, continuous, stochastic) these models cannot be studied using formal analytical tools, and thus the analysis and exploration of simulated trajectories also require complex workflows to guide the exploration the parameter spaces associate to these models [29]. For this reason, distributed workflows to perform parallel optimization via simulation and model exploration are critical tools to exploit the full potential of simulations [28, 30]. Model exploration workflows are required to efficiently fit parameters for which there are no available experimental measurements [1, 27] and also to explore complex and vast parameter spaces and to optimize user desirable goals, such as the space of optimal treatment strategies for a given cancer model. Optimization methods such as evolutionary algorithms have proven their usefulness in such studies, for fitting unknown parameters [1], as well as, high-throughput hypotheses testing in cancer research [27].

In previous work, Letort et al (2019) developed the multi-scale model of 3T3 fibroblast spheroids that integrates the Cell Fate Boolean network [5] inside individual cell-agents. The Boolean network rules the phenotype of the cells (e.g. proliferation, apoptosis) based on the environmental conditions (e.g. drugs presence, oxygen concentration). The authors used the model to investigate the tumor response to different regimes with tumor necrosis factor supplies (TNF) and reported complex behaviors in the simulated conditions. While a set of values of pulse period, pulse duration and TNF concentration that was optimal to reduce the number of alive tumor cells, different sets of values turned the cells resistant to TNF [22]. The effects of TNF in the Boolean model reported by Calzone et al. (2010) are multifaceted: TNF triggers cells to go from a Naive to a Proliferative state, but also to commits cells to Necrosis and Apoptosis. Once the cells are committed to either Survival, Necrosis or Apoptosis, they cannot go back, causing resistance due to phenotypic variability. Interestingly, it has been described that prolonged TNF exposure causes the cells resistant to the effect of the cytokine [21].

In the present work, we use an extended hybrid multi-scale model to perform a deep exploration of the parameter space of effective treatment strategies based on TNF pulses with the aim of unravelling the mechanistic details behind the complex emergent dynamics of the TNF pulses in *in-silico* experiments; and also to guide the optimization of effective treatments. We extended the multi-scale model of 3T3 fibroblast spheroid by integrating an explicit kinetic description of the TNF-receptor dynamics, based on the molecular biology of the TNF receptor [7, 23, 31]. Furthermore, we couple the TNF-receptor kinetic model with the cancer cell Boolean model from [5] to simulate the downstream propagation of the signal that induced the binding of the TNF. To explore the parameter space of effective treatments in different scenarios and conditions, we have developed an HPC-optimized model exploration workflow based on EMEWS [28]. Our workflow includes two previously used model exploration strategies, Sweep Search, Genetic Algorithm [27, 1], together with a new approach, named the Covariance Matrix Adaptation Evolutionary Strategy which has been shown to exhibit good convergence in global optimization problems with continuous variables [14].

We applied our framework to characterize the space of effective treatments in different experimental scenarios, by simulating the treatment outcome with our multi-scale model of a tumor growth. We first studied the effect of the cells spatial distribution in the values of the treatment parameters by optimizing the supply strategies in 2D monolayers and 3D spheroids of different sizes. We found that our model exploration workflow is able to find non-trivial *in-silico* drug scheduling strategies that minimize the tumor below 1% of its initial size while avoiding the emergence of resistant cells. Our results also show that effective treatment strategies can be found in the two different cell geometries studied. We also found that the parameter spaces of effective treatments for the 2D monolayer and 3D spheroid exhibit different parameters distributions of the parameters. We later study the robustness of the effective treatments when heterogeneous populations of cells are considered. Specifically, we model population heterogeneity by introducing different levels of cell-based variability into the kinetic parameters of the TNF receptor models. The parameters’ variability aims to mimic population-level variability in the kinetic parameters of the receptor, as well as, different levels of expression in the receptor, among different cells. We found that effective treatment strategies are robust to a low level of variability, nonetheless, whereas with a high level of variability, those treatment strategies optimized for populations with no variability cannot reduce tumor growth. However, when the treatments are optimized directly on an heterogeneous population we observed that the optimization algorithms can retrieve effective treatment.

Altogether, we found that our model exploration workflow can find effective treatments in all the studied conditions, showing the multi-scale simulations and model exploration are promising tools for *in-silico* exploring treatment strategies. Finally, our results also show that cells’ spatial geometry, as well as, population variability should be considered when optimizing treatment strategies in order to find robust parameter sets. In future work we plan to extend these results by studding other experimental setups as well as different cancer models.

## Materials and Methods

### Hybrid multi-scale model of cancer cells with signaling

Herein, we present a multi-scale model of tumor growth that considers at the individual cell level the dynamics of the tumor necrosis factor (TNF) receptor and its downstream effect using a hybrid approach (Figure 1). Our model was implemented using PhysiCell [12] together with the PhysiBoSS add-on (Ponce-de-Leon et al., *in preparation*). The microenviroment is simulated both in the 2D and 3D domains, and it accounts for the presence of oxygen and of the cytokine tumor necrosis factor (TNF). On the other hand, cell are simulated as individual agents which include intracellular submodels that account for the cell cycle, the different death models (i.e. necrosis, apoptosis), a model for TNF receptor dynamics, and a gene regulatory network.

**Figure 1:**
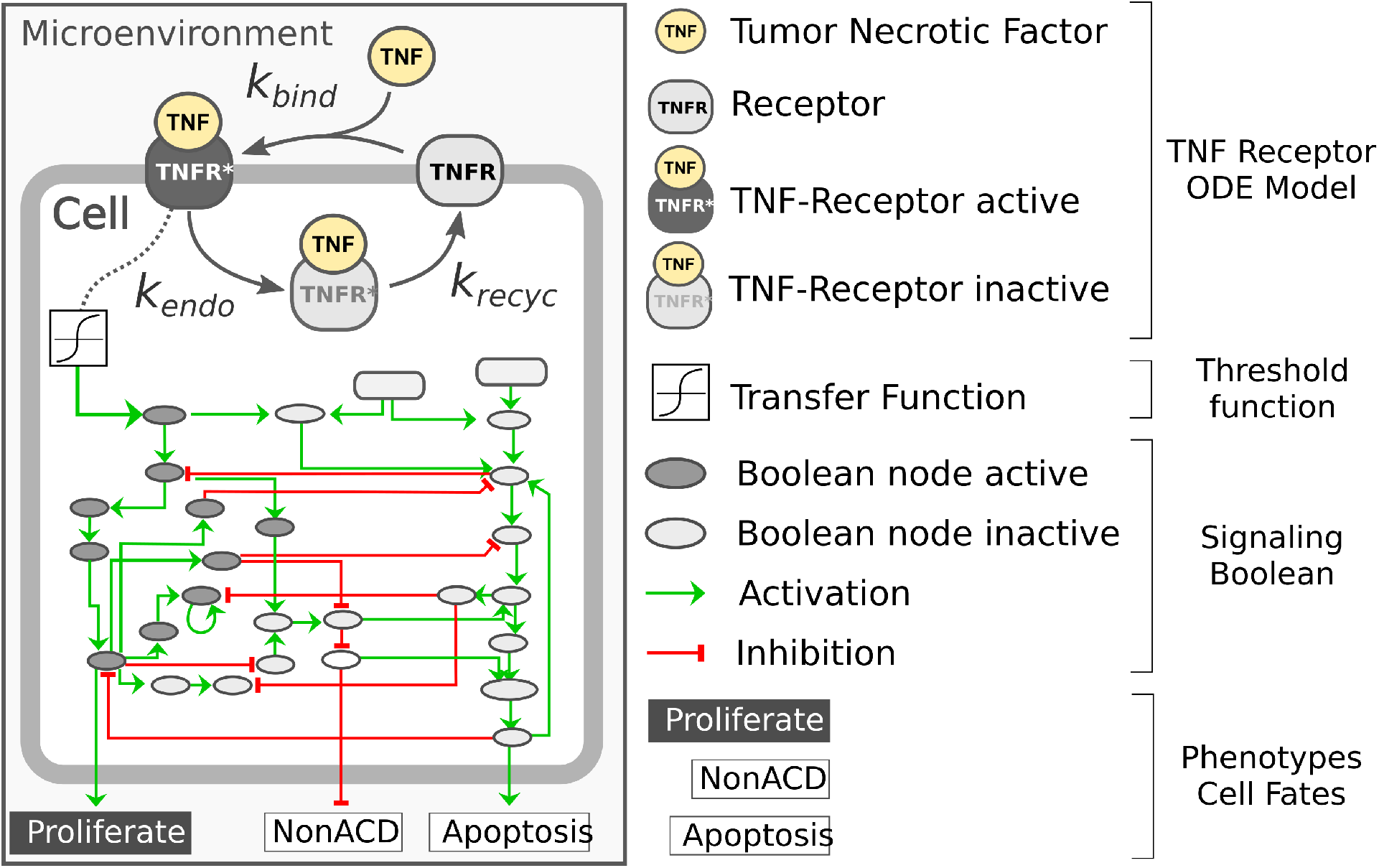
Diagram representing the intracellular submodels of the multi-scale model of a tumor growth. Each individual cell agents has a kinetic model of the TNF receptor dynamics that is connected to the microenvironment through the presence of surrounding TNF and that is coupled to the Boolean network through a transfer function. The Boolean network has three readout nodes (Proliferation, NonACD and Apoptosis) which rule the fate of the cell agente

For the cell cycle, we use PhysiCell *live cell cycle* with a doubling time of 22*h* and for the death models, we used PhysiCell standard ones with default parameters. The binding of the TNF to its receptor is modeled using mass-action kinetics in which TNF binds to a cell receptor TNFR at a given rate *k_bind_*; the complex TNF-TNFR is internalized at a rate of *k_endo_* where the TNF is degraded and the receptor recycled at a rate of *k_recycle_* (see Supplementary Figure 1).

TNF-receptor submodel was developed based on the known molecular biology of the molecular system [7, 23, 31]. The equation below describes the submodel for the TNF receptor dynamics.

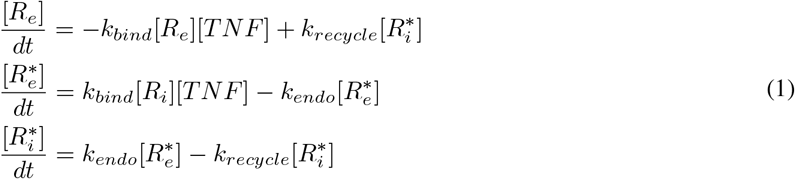

where [*R*], [*TNF*] and [*R*\*] are the concentrations of the receptor, the TNF, and the TNF-TNFR complex, respectively. Furthermore, the TNF-TNFR complex [*R*\*] can be in two states, in the cell membrane 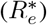 or internalized 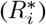.

The gene regulatory model used is an extended version of the Boolean network (BN) reported in Calzone et al (2010) and is simulated using the MaBoSS algorithm. The BN is coupled to the agent in two different ways (see Supplementary Figure 2). The BN has an input node that represents the presence of *TNF* and is coupled to the amount of active TNF-TNFR complex 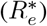 through a transfer function that converts the continuous value of 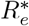 into a Boolean one. Additionally, the BN has three mutually exclusive output nodes that represent three alternative cell fates which include, *Proliferation, Apoptosis*, and *NonAD* (non-apoptotic death or necrosis). The fate or phenotype of each cell agent is ruled by the current state of the fate nodes of its internal regulatory network. For instance, if the *Proliferation* node is active the cell will grow and divide, whereas if the *Apoptosis* becomes active the cell agent will commit to apoptosis (see Supplementary Figure 2).

### Model exploration framework

#### Workflow overview

The parallel simulation framework that is used in our evaluation is a workflow that follows the Extreme-scale Model Exploration with Swift (EMEWS) paradigm, it uses the *spheroid_TNF_v2* as an example model, and is publicly available in our online repository.^1^ An overview of our model exploration workflow is shown in Figure 2. We have integrated three different search strategies: (*i*) a Sweep search approach that evaluates a predetermined set of candidate parameters (generated from uniform sampling, or a regular grid), (*ii*) a Covariance Matrix Adaptation Evolutionary Strategy (CMA-ES), and (*iii*) a Genetic Algorithm (GA). For the cases of CMA-ES and GA we use the available implementations provided by the DEAP package (version 1.3.1) a mature and widely used package for evolutionary optimization [10]. Using EMEWS queues, multi-scale simulation instances are configured with the specific parameter values that correspond to the points that each exploration procedure targets, and are then submitted for parallel execution in an HPC environment.

**Figure 2:**
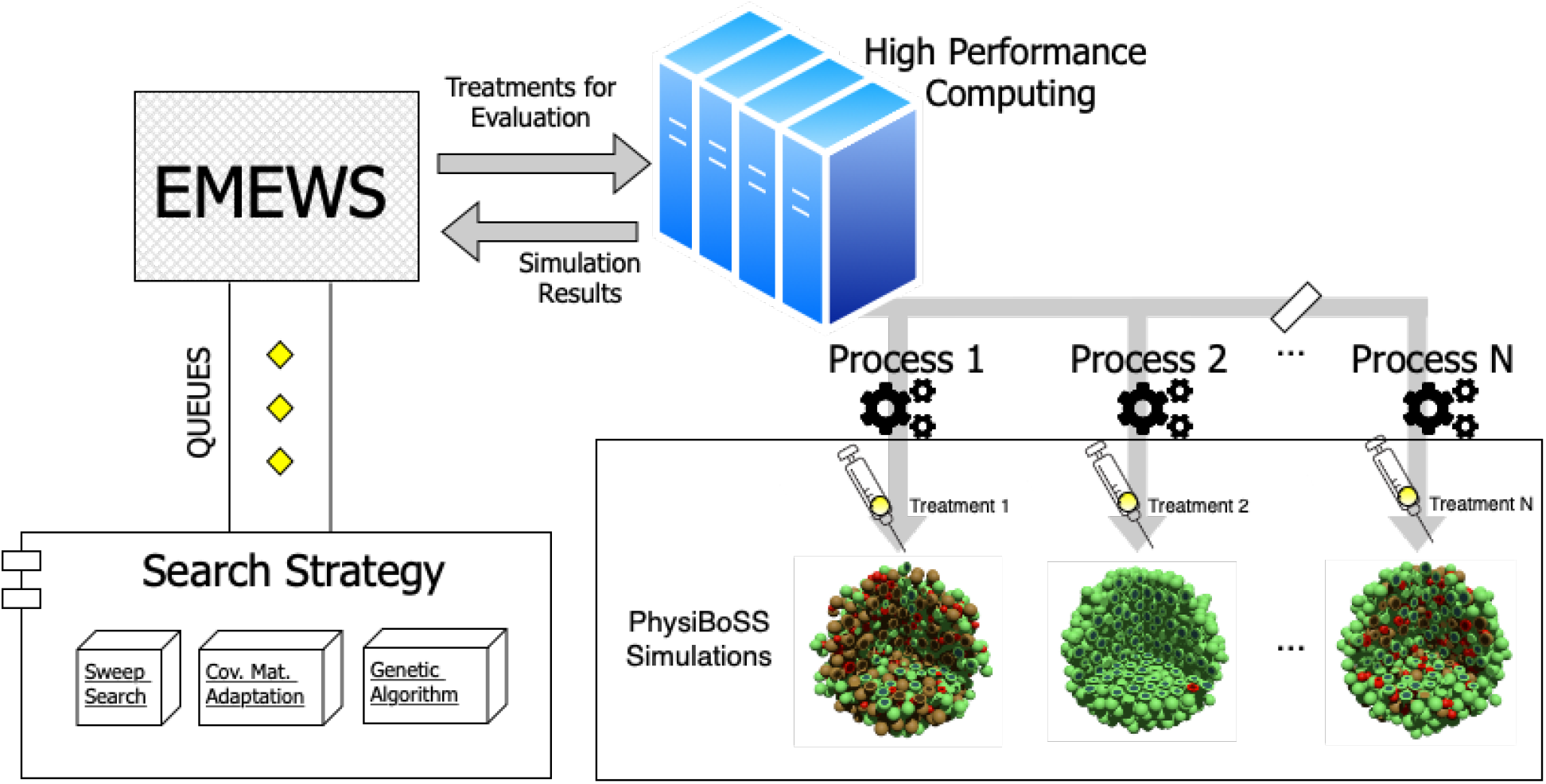
Workflow overview. The diagram depicts the structure of the model exploration workflow. EMEWS communicates to the different search strategies using a queue system. The search strategy generates candidate parameters, and the treatments to be evaluated via PhysiBoSS simulations are distributed as parallel jobs to the HPC infrastructure. Upon completion, the outcomes of the simulation are returned to EMEWS which, in the case of the GA and CMA-ES, sends the fitness of the evaluated parameters so the algorithm can update its internal state and generate new candidate parameters.

The number of each ‘batch’ of points is relative to the number of computational nodes that are available in the HPC. The multi-scale simulator incorporates PhysiCell (v1.7) [12] together with the PhysiBoSS 2.0 extension (Ponce-de-Leon et al, in preparation), which is an add-on version of PhysiBoSS. We merged our mass-action kinetics model, as explained in Section Hybrid multi-scale model of cancer cells with signaling, with the multi-scale model proposed by Letort at al (2019) that used the Boolean model from Calzone et al. [5]. Finally, the simulation results are returned, the points are evaluated according to the performance of the particular drug treatment, and the workflow iterates over the next ‘batch’ of points. Each point is a three-dimensional vector that configures the following simulation parameters: i) the duration of the TNF pulse; ii) the TNF pulse period; and iii) the concentration of TNF. The exploration space has ranges of from 5 to 800 minutes for the pulse period, from 5 to 200 minutes for the pulse duration, and from 0.001 to 1 ng/L for the TNF concentration.

We check the number of alive tumor cells at the last time-point of each simulation to evaluate the results of each particular treatment. Note that, to ensure that the characterization is robust and not a subject of extreme randomness accruing from inherent PhysiBoSS stochasticity, we perform 3 replicate simulations with the same configuration parameters, using a different seed to initialize the random number generator, and calculate the average value of the final alive tumor cells count over the replicates as the final score. We now proceed to describe the different search methods we employ in detail.

#### Sweep search

The Sweep search comprises a simple exhaustive approach that requires the user to specify a predetermined number of points to be evaluated. Our code offers a points generating script, which can be configured to choose among different distributions. That is, points can either be selected to belong on a grid, with equal distances between each point along the dimensions, or a second option is to select random points by sampling particular probability distributions. For the purposes of the experimentation presented in this paper, we have implemented the uniform distribution point selection, though this can be easily configured to employ other types, e.g. Gaussian, Beta, etc.

#### Covariance Matrix Adaptation Evolutionary Strategy

Covariance Matrix Adaptation Evolutionary Strategy (CMA-ES) is a stochastic, derivative-free method for numerical optimization for black-box optimization functions [14]. This method requires as input a set of points, a *σ* value that controls the range of exploration, a covariance matrix *C* that is used to guide the search, the number of points population to execute the algorithm upon, and a total number of iteration or stop criteria. In a nutshell, CMA-ES generates an initial population sampling from a multivariate normal distribution, evaluates each generated point, and then calculates mutation steps of the best points to form the mutation distribution. By this, a new population of points is generated and evaluated, and the iterative process continues up to a user-defined number of times. Note that, for every update of the mutation distribution in each algorithm iteration, all past paths from previous iterations are also taken into account, and the most favourable points are granted larger probability for being selected by the evolutionary strategy. This way, the length of each mutation step can be adapted to be longer in cases of greater fitness score improvement, or shorter for the opposite case.

#### Genetic Algorithm

Genetic Algorithm (GA) is a widely known and tested metaheuristic approach, also belonging to the family of evolutionary strategies algorithms, that mimics the evolution principles of biological organisms and operates directly on the values of points [15, 36]. Similarly to CMA-ES, an initial population of points is generated and evaluated, and then, following an iterative approach, a series of genetic operators are applied to each of them, in order to produce the next, evolved, the population of points. More specifically, the GA applies the selection, crossover, and mutation operators. Typically, the first operator selects the evaluated individuals in a weighted manner, so that the ones with better fitness scores have an increased probability of being selected to proceed to the next generation, compared to fewer fit points. Then, the crossover mixes the point values in a principle similar to that of the gene propagation from parents to offspring as it happens in organisms. Finally, the mutation operator changes a point value (e.g. one of its dimensions) with a small probability, similarly to the process that has been observed in DNA sequences. The main idea is that combinations of points with good fitness scores would lead to even better ones, especially if the search domain is smooth. However, since the algorithm takes into account only the previously observed fitness scores, without having any other domain-specific knowledge, as a consequence it may happen that its search is constrained around locally optimal points, never managing to reach the global optimal ones. In spite of this shortcomings, GAs have been shown to work very well for non-smooth search spaces as well [35, 8].

## Results

### Multi-scale simulations and model exploration setup

Herein, we use a multi-scale model of tumor growth to investigate different treatment strategies. The model, which is also used in previous work [1], simulates the dynamics of a population of cancer cells growing under different drug treatment conditions. A treatment strategy consists in the supply of periodic pulses of the cytokine tumor necrosis factor (TNF), with fixed duration and concentration (see Sec. Materials and Methods). At the molecular level, when the TNF binds to the cell’s receptor TNFR forming a complex, and the TNF-TNFR complex concentration reaches a given threshold, the signal is propagated through the Boolean regulatory network inducing cell death. However, if the stimulus is sustained for a longer period of time, cells activate NFkB node and the Survival node, becoming resistant to the death induced by the TNF. For this reason, optimal treatments should expose the cell for a sufficient time to induce death but not too much as to become resistant to it [22].

To explore the parameter space associated with the treatment, we have extended our model exploration workflow based on EMEWS [1]. In each *in-silico* experiment, we simulate the growth of a population of cancer cells for 4640 minutes (i.e. three days) subject to a given treatment strategy. To account for the inherent stochasticity of the model, each simulation is always run in three replicates and the average behavior is considered (see Sec. Materials and Methods). We evaluate the effect of the treatment strategies by analyzing the total number of alive cells at the end of the simulations, relative to the initial population size, and use these values as the score, or *cost function* associated to a treatment strategy. We define as a *effective treatments* those strategies that reduce the number of the alive cell below 1% of its initial numbers, in the three replicates. Based on this definition, we investigate the parameter space of the *effective treatments* in two different spatial arrangements of cells: a mono-layer disc of radius 100*μ*m (151 cells), and a 3D spheroid of radius 100*μ*m (1173 cells).

### Effective treatment parameters differs for 2D and 3D cell arrangements

To investigate the structure of the parameter space of the *effective treatments*, we perform a uniform sampling of 10, 000 candidate sets of parameters corresponding to different treatment strategies. We use these sets of parameters (sweep search) as inputs for multi-scale simulations and we evaluated each treatment effect on the growth of the cancer cells in the 2D and 3D arrangements. From the 10, 0000 evaluated parameter sets, the results show that 113 strategies are *effective treatments* for the 2D setup, whereas in the 3D case, only 11 strategies are *effective treatments* (see *Supplementary Table 1*). This indicates that the region containing the *effective treatments* for the 3D spheroid is more constrained than in the 2D disc arrangement. Interestingly, if we constrain the definition of *effective treatments* to kill all cancer cells at the end of the simulation and in the three replicates, only 8 sets of parameters can reach the goal for the 2D cases whereas no effective parameter sets are found for the 3D setup (see *Supplementary Table 1* and Supplementary Figure 3).

We compared the distributions and summary statistics of the parameters of the *effective treatments* in the two arrangements (Figure 3). The comparison shows that the distribution values for the evaluated parameters indicate that the *effective treatments* of the 3D arrangement are notably more constrained than those that work in the 2D arrangement, in particular regarding the concentration of TNF and the *Pulse duration*. In general, the *effective treatments* in 2D arrangements exhibit bigger values and larger ranges in the three parameters (Table 1).

**Figure 3:**
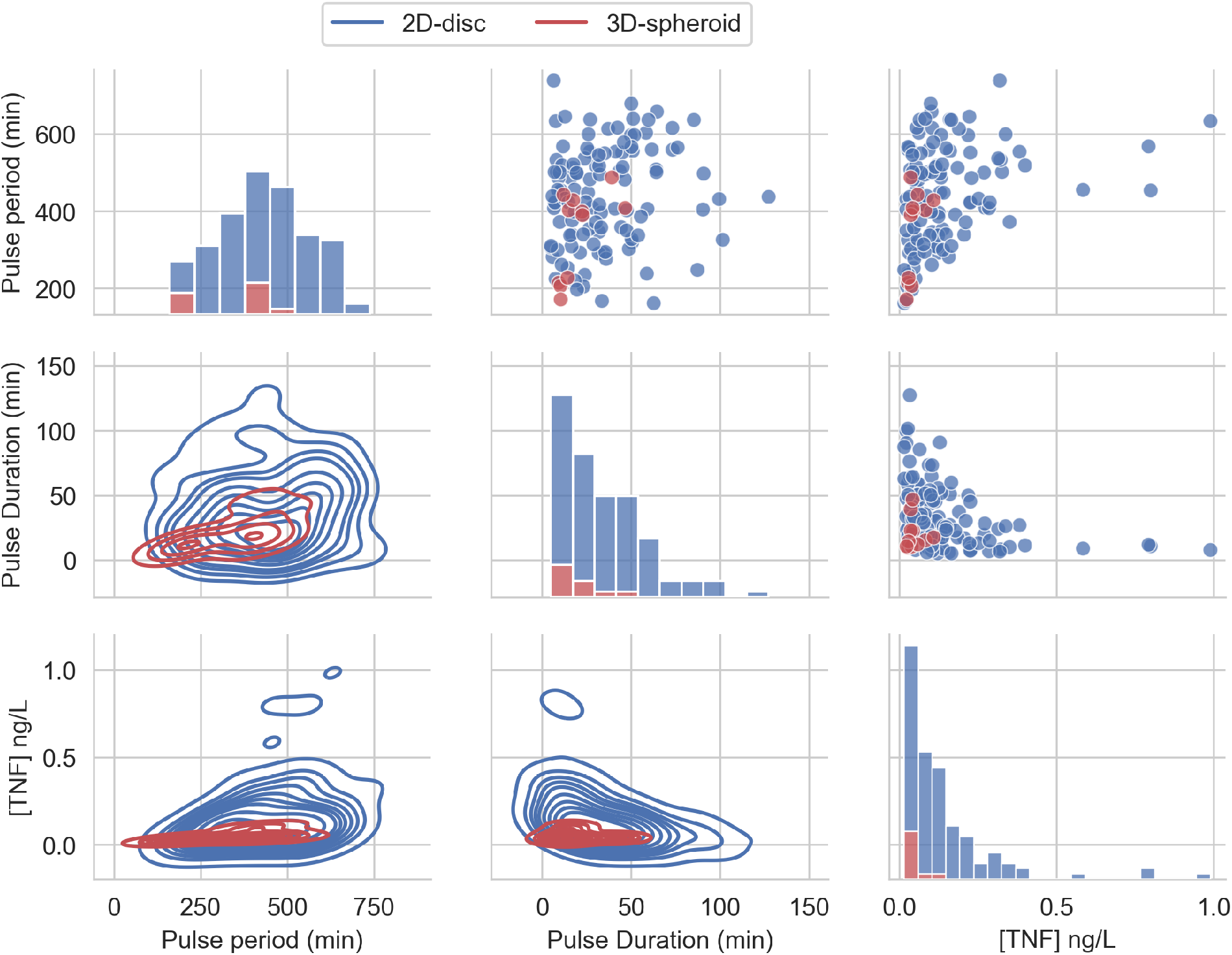
Effective TNF treatment parameters distribution from uniform random sampling. The figure depicts the distribution of the three parameters of the pulse treatment, sampled from a uniform distribution and filtered to belong to the feasible region, i.e. that can reduce the tumor size below 1% of its initial size.

**Table 1:** Summary statistics for the parameters from the *effective treatments* found by sampling 10.000 random candidates.

| Parameter | Layout | Mean | Std | Min | Median | Max |
| --- | --- | --- | --- | --- | --- | --- |
| *Pulse Duration (min)* | 2D | 34.66 | 24.54 | 5.00 | 30.17 | 127.21 |
|  | 3D | 19.97 | 12.31 | 9.37 | 14.99 | 46.66 |
| *Pulse period (min)* | 2D | 440.83 | 128.53 | 161.44 | 438.86 | 738.09 |
|  | 3D | 343.28 | 113.64 | 171.28 | 398.79 | 487.19 |
| *[TNF] (ng/L)* | 2D | 0.14 | 0.16 | 0.02 | 0.10 | 0.99 |
|  | 3D | 0.05 | 0.03 | 0.02 | 0.04 | 0.11 |

We also found that some parameters’ combinations exhibit correlations (see Supplementary Figure 4). On the one hand, we observe that the *Pulse period* positively correlates with [*TNF*] in the 2D and 3D but only correlates with *Pulse duration* in the 2D case. On the other hand, [*TNF*] shows a negative correlation with *Pulse duration* only for the 2D case. Altogether, these correlations indicate that the treatment parameters can compensate each other: a shorter pulse might be as effective as a longer one if it carries more [*TNF*].

**Figure 4:**
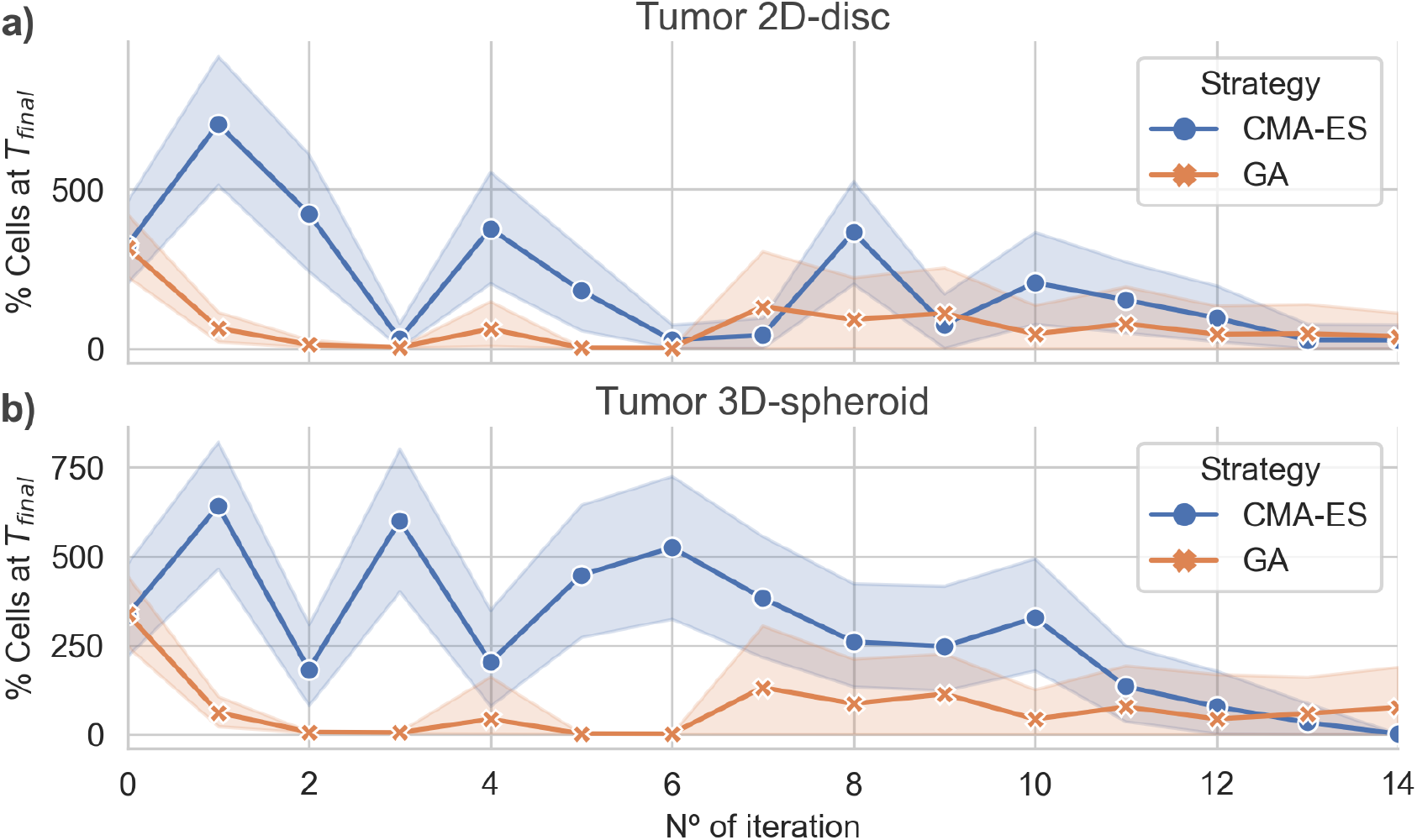
Algorithmic convergence for the optimization of treatment parameters. The figure depicts the average % of alive cells at the end of for each iteration step of the CMA-ES and GA optimization algorithms. Panels a) and b) show the convergence of the algorithms for simulations considering a population of cancer cells arranged in a 2D disc and a 3D spheroid, respectively.

### Optimal treatment parameters differs for 2D and 3D cell arrangements

To further investigate the structure of the parameter space of the *effective treatments,* we conducted an optimization via simulation to find the set of treatment parameters that minimizes the number of alive cells at the end of the application of the treatment. We performed the parameter optimization in both cell arrangements (i.e. 2D disc and 3D spheroid of radius 100 *μ*m) focusing on the same parameters as in the previous section Effective treatment parameters differs for 2D and 3D cell arrangements: *Pulse duration, Pulse period* and [*TNF*]. These optimizations were run using two evolutionary algorithms: Covariance Matrix Adaptation Evolutionary Strategy (CMA-ES) and Genetic Algorithm (GA) (see Sec. Materials and Methods for details).

The results showed that both algorithms converge to optimal (leaving no remaining cells) or near-optimal sets of parameters for the 2D and 3D case, respectively (Figure 4). Interestingly, in both cases the GA found *effective treatments* after 2 iterations, whereas the CMA-ES only does after 15 iterations. Nonetheless, both algorithms are capable of finding *effective treatments*. In addition, the CMA-ES algorithm also estimated a multivariate normal distribution for the region of *effective treatments* parameters that is updated at each iteration. In the last iteration, the population sampled by the CMA-ES showed a very low variance (Figure 4) in the 2D and the 3D arrangements, indicating that the estimated distribution captures, at least, part of the structure of the parameters associated with the *effective treatments*.

We compared the parameter sets for the *effective treatments* predicted by both algorithms and uncovered that each one converges to different regions of the parameter space (see Supplementary Figure 5). The CMA-ES found *effective treatments* parameter distributions that are different for the 2D and 3D. In both cases, the parameter ranges were narrower than those found in uniform sampling. Moreover, the CMA-ES converged to distributions for the *Pulse period* that were different for the 2D and the 3D arrangement. In addition, the parameters corresponding to *effective treatments* found by the GA were more scattered showing wider ranges of values. In both cases, the *Pulse duration* seemed to be the less critical or an unconstrained parameter.

We analyzed the time course of the total number of alive, apoptotic, and necrotic cells for one of the optimal treatments found. Furthermore, for this simulation, we also checked the internal state of TNF receptor model (i.e. the values for R_e_, 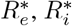 variables) at each time step of the simulation, and averaged these values over all the cells to investigate the coarse grain dynamics of the submodel. Figure 5a shows how the number of alive cells periodically drops until it reaches zero, as a consequence of the *TNF* pulses. When analyzing the average dynamics of the TNFR receptor model in *effective treatments* we found that the TNF pulse needs to trigger the activation of the receptor of 50% of the population for short periods of time to be effective (5b). If the average number of cells that got activated is lower than this threshold, then the rate of cells entering necrosis will be lower than the population growth rate, and therefore the number of alive cancer cells will steadily grow. On the contrary, if the average number of cells that got activated is above this threshold for extended periods of time, many cells will become resistant to the treatment. We found similar results for the case of the 3D spheroid (data not shown).

**Figure 5:**
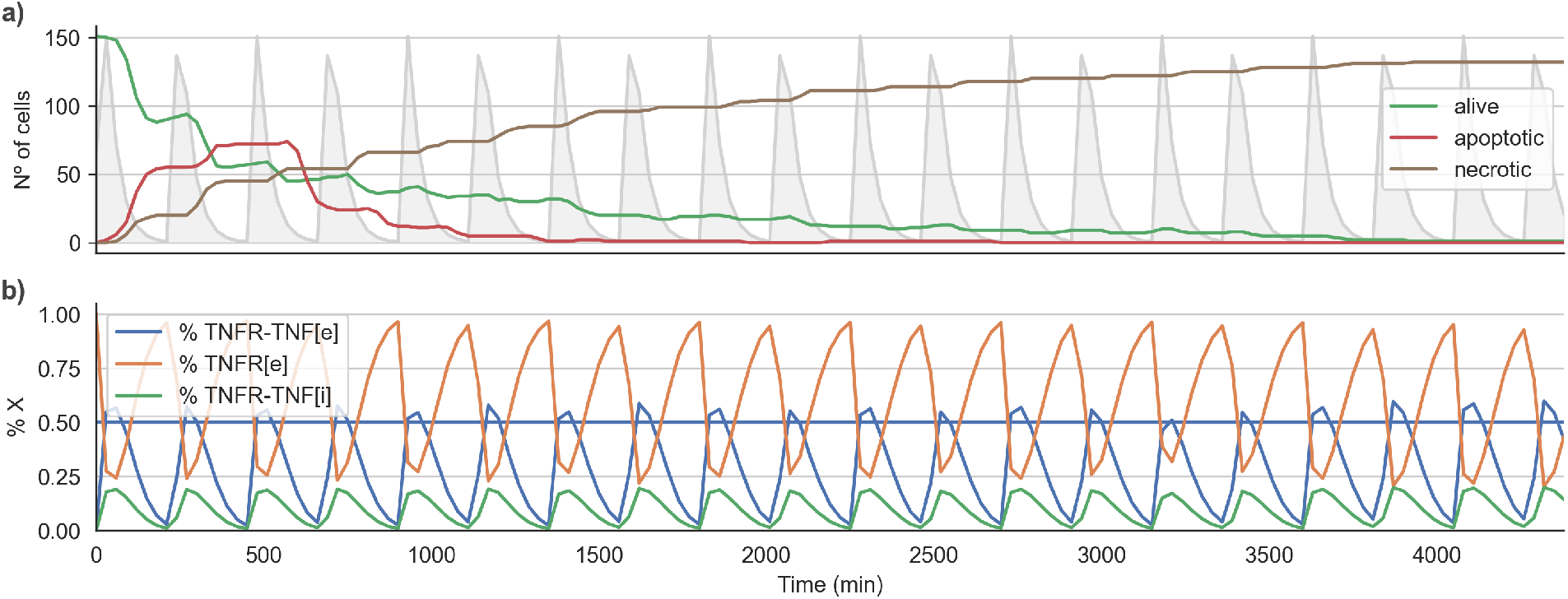
Time course for an *effective treatment* in 2D cell arrangement. Panel a) shows the time course for the number of Alive, Apoptotic an Necrotic cells. The lightgrey curve shows the *TNF* pulses. Panel b) shows the average state of the TNFR receptor model across all the alive cells. The horizontal line indicates the threshold at which the signal induced by the binding of the TNF propagates downstream to the Boolean network.

### Robustness analysis of the *effective treatments* in heterogeneous populations

In the previous section, we showed that *effective treatments* can be found for 2D and 3D cell arrangements, using either the CMA-ES or the GA algorithms. Nonetheless, those treatments were optimized on monoclonal or homogeneous tumors, i.e. population of cells with identical parameters. In this section, we study the robustness of *effective treatments* by studying heterogeneous populations of cells. We forced the population heterogeneity by introducing variability into the three kinetic parameters of the TNF receptor, i.e. the TNFR binding rate, the TNFR endocytosis rate and the TNFR recycling rate. The variability is applied by considering a normal distribution centered in each parameter’s default value and with a standard deviation, which is the control parameter, that varies from zero (homogeneous population) to one (almost uniformly distributed random parameters). Then, when the population is initialized, the kinetic parameters of each cell are sampled from the corresponding distribution.

To evaluate the robustness of the *effective treatments* in heterogeneous populations, we considered the top 30 *effective treatments* parameter sets that had no final tumor cells in any of their replicates for the 2D and for the 3D cell arrangements. Then, for each set of parameters we run the simulations with different levels of variability from 0 to 1. As expected, we observed that for low values of the variability control parameter (< 0.2) most of the evaluated *effective treatments* still can reduce the initial tumor size to the 1% of the initial size. However, for higher variability values (> 0.2) most of the evaluated treatments could not reduce the tumor size below its initial size (Figure 6). This indicates that as the level of variability on the TNFR receptor kinetic parameters increases, the effectiveness of the treatments dramatically decreases (Figure 6).

**Figure 6:**
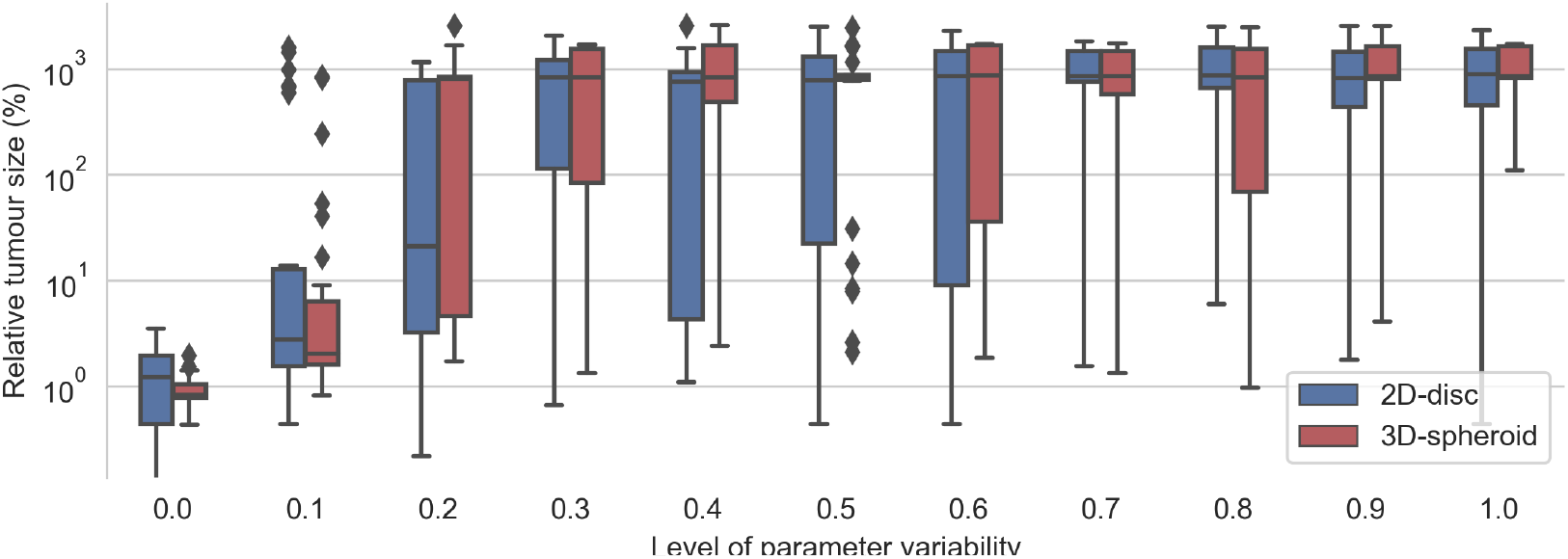
Evaluation of the top 30 best effective treatments in heterogeneous populations of cancer cells with different degree of variability in the kinetic parameters of the TNFR. For a given value kinetic parameters’ variability (x-axis), each pair of boxes depicts the distribution % of alive cells at the end of the simulations obtaining after evaluating the top 30 best effective treatments founded when zero variability was considered.

Interestingly, the critical value at which most of the *effective treatments* were no longer effective is different for the 2D and the 3D cell arrangements. In the case of the 2D disc, the critical value is close to 0.25 whereas in the 3D spheroid cases this value is around 0.15 (Figure 6). When the variability value is above this threshold, some of the sampled kinetic parameters make the cell insensitive to the treatment and thus it can grow even in the presence of TNF. The differences in the critical threshold were possibly due to the different number of initial cells considered in the 2D and the 3D cell arrangements. To assess this, we tested the effect of the radius size of the 2D simulations in this robustness analysis.

We evaluated radius sizes of 50, 275 and 500 μm which corresponds to initial tumor sizes of 37 cells, 1069 and 3559, respectively. We tested three levels of variability with each radius size in three replicates and averaged their results. As already discussed, we can observe that the more variability, the worse the outcome (see Supplementary Figure 6). Interestingly, we did not see a clear correlation between the radius length and the decrease of the effectiveness of the treatments, with the 50 *μ*m being the one with worse outcomes with the higher variability level.

### Optimization via simulation can find effective treatments in heterogeneous populations of cancer cells

To evaluate the performance of model exploration workflow in more complex scenarios, we investigated the optimization of treatment strategies in tumors with different levels of heterogeneity. This use case was considered as a way to evaluate what would happen when using this methodology in a less ideal situation as it can be the drug screens in cell lines or tumors with heterogeneous non-clonal cells. For this purpose, we conducted the optimization via simulation to find effective treatments in two conditions with different degrees of variability in the kinetic parameters.

At first, we set the variability value on the kinetic parameters of the TNFR receptor model to 0.25 and then run the GA and CMA-ES to find treatments that minimize the total number of alive cells. We found that with this level of variability neither of the algorithms were able to converge (see Supplementary Figure 7). Nonetheless, this did not prevent the algorithms to find *effective treatments* for both arrangements: the 2D disc and the 3D spheroid. While for the 2D disc several candidate sets of parameters were found, only a few candidate *effective treatments* were found by the CMA-ES for the 3D spheroid.

Interestingly, for the 2D cell arrangement, the CMA-ES algorithm was able to find two optimal treatment strategies, i.e. a set of parameters for the *TNF* pulse that kills every tumor cell in all three replicates. These two sets of treatment parameters are similar to those *effective treatments* that worked when variability was not introduced (see previous section). Figure 7a shows the time course for the *effective treatments* optimized in the 2D cell arrangement. The plot shows how the number of alive cells steadily decreases until zero (Figure 7b). The Figure also shows the average internal dynamic of the receptor model exhibiting some noisy behavior due to the heterogeneity present in the population.

**Figure 7:**
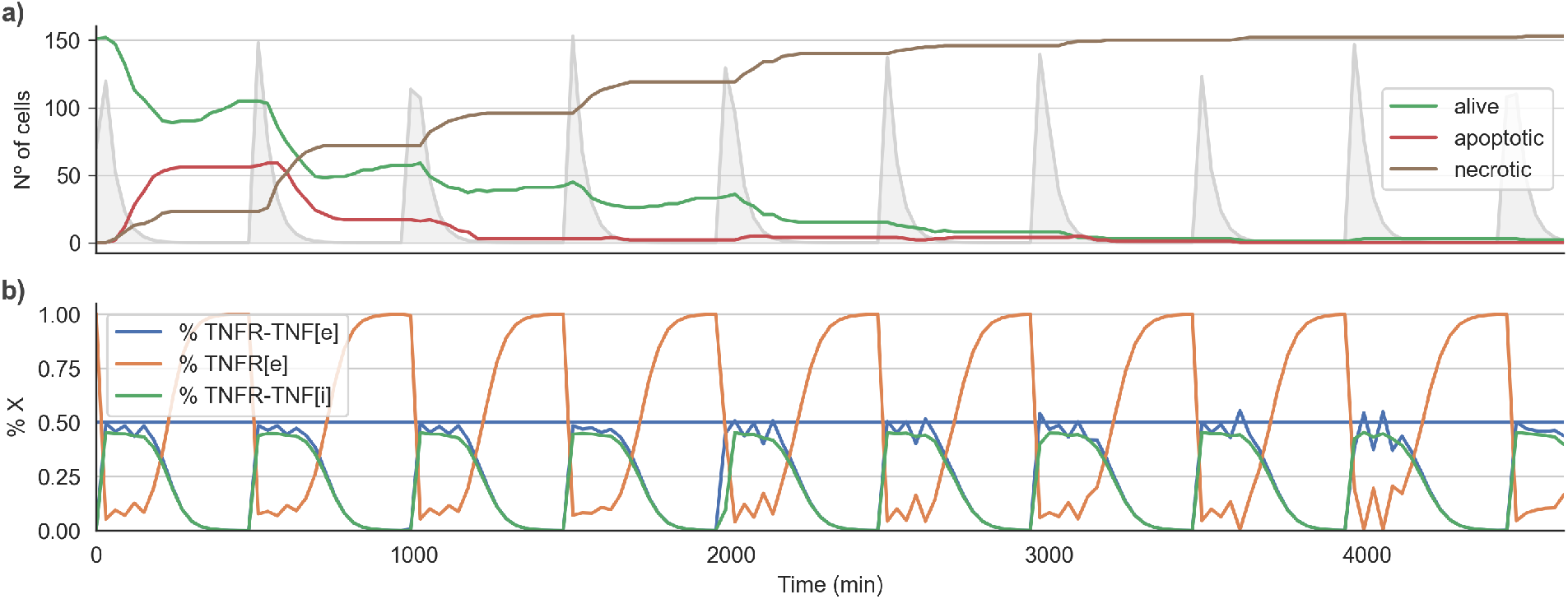
Time course for an *effective treatment* in the 2D cell arrangement with a variability of. 0.25. Panel a) shows the time course for the number of Alive, Apoptotic an Necrotic cells. The lightgrey curve shows the *TNF* pulses. Panel b) shows the average state of the TNFR receptor model across all the alive cells. The horizontal line indicates the threshold at which the signal induced by the binding of the TNF propagates downstream to the Boolean network.

We also performed a similar experiment with a higher variability value (0.50) on the kinetic parameters of the TNFR receptor model. We ran the treatment optimization using the GA and CMA-ES and found that, as expected, with this level of variability the convergence of both algorithms was even worse than for the case of 0.25 (see Supplementary Figure 8). Furthermore, the algorithms were only able to find *effective treatments* for the 2D disc arrangement. Nevertheless, the number of different parameter sets found were fewer than in the previous scenario with a variability value of 0.25, as expected for a more complex landscape. For the 3D spheroid case, the best treatment reduced the initial tumor size to 1.05% of its initial size when averaged over the three replicates. If we relax the definition of *effective treatment* to a threshold of 2% and compare the distribution of the effective parameters between the cases with variability set to 0.25 and 0.50, we found that the ranges of values were wider in the second case. Altogether, the results presented in this section indicate that the higher the variability in the population the harder is to find *effective treatments*. Nonetheless, the results also show that even with high values of parameters variability it is still possible to find very effective treatment strategies.

## Discussion

In this work, we used a hybrid multi-scale model that merges a mass-action kinetics model of the TNF receptor with a cancer cell Boolean model of different signaling pathways. Moreover, these models are embedded in an agent-based framework that allows considering populations of cells in a defined microenvironment. By performing a model exploration, we have shown that the *effective treatments* parameter can be found in different cells’ geometries including 2D monolayers and 3D spheroids. Furthermore, by performing a uniform random sampling of the *effective treatment* spaces we found that the parameters for 2D and 3D arrangements exhibit different distributions. These differences are more pronounced in the case of the TNF concentration as well as the Pulse duration, where the effective treatments for the 3D spheroid case are notably more constrained than those of found for the 2D disc. We hypothesize that the 3D configuration imposes spatial constraints in the diffusion of the TNF which restraint the space of vales for the candidate *effective treatment*.

We also found that some parameters’ combinations exhibit correlations indicating that one parameter change can be compensated by adjusting another one. For instance, we observe that the Pulse period positively correlates with [TNF] in the 2D and 3D. This means that increasing the period between pulses can be compensated by an increase in the pulse concentration. Interestingly, the correlation between these two parameters is stronger in the 3D spheroid than in the 2D monolayer, which also shows how the former case is more constrained than the latter. For the 3D spheroid we also found a strong positive correlation between pulse period and its duration, showing that these two parameters can also compensate each other. Nonetheless, the correlation between these two parameters is very low and does not show statistical significance in the 2D monolayer. Finally, we found a negative correlation between the pulse concentration and its duration in the 2D monolayer showing that in these cases an increase in the concentration of the pulse can be compensated by a reduction of its duration. Altogether these results indicate that the structure of parameters spaces of *effective treatment* is depends on the spacial cell distribution and therefore treatment strategies that work in a 2D monolayer may not work in a 3D spheroid.

We later investigate the optimization of treatment using two different evolutionary algorithms, GA and CMA-ES. Our results showed that both algorithms can quickly converge to *effective treatments* but while the GA can find candidates in the first iteration the CMA-ES converge to a more robust region of the parameter space. Furthermore, the CMA-ES also finds a statistical distribution for the region *effective treatments.* Strikingly, both algorithms converge to slightly different regions of the parameter space in in both the 2D and the 3D arrangements. This suggests that the fitness landscape of *effective treatments* is very rough exhibiting several valleys of *effective treatments* regions.

In order to unravel the molecular mechanism behaving the effective treatments, we analyzed the coarse-grained dynamics of the TNF receptor models for working and none working treatment strategies. The non-working strategies can be grouped into two classes. On one hand, there are those sets of treatment parameters that do not allow the receptor to reach the threshold needed to propagate the signal downstream the Boolean model. On the other hand, there are treatment strategies that keep the activation of the receptor for long periods enough to induce cell resistance. In this context, if we define drug resistance as the inability of a cell to respond to a given drug, we find that resistance can come from two aspects: from the dynamics of the Boolean model in response to TNF and from the characteristics of the TNF receptors. The effects of TNF in this Boolean model reported by Calzone et al (2010) are multifaceted: TNF triggers cells to go from a Naive to a Survival state, but also to commit cells to Necrosis and Apoptosis [5]. Once the cells are committed to either Survival, Necrosis or Apoptosis, they cannot go back, causing a resistance that can be dues to phenotypic variability. As this model was studied using a stochastic Boolean simulator [33], it was possible to capture its dynamics and see that these commitments were not equally fast, or even, that there was a window of activation that allowed to control the commitment to Survival and commit the cells to Necrosis [22].

In addition to the Boolean model, our hybrid model also has a mass-action kinetics model that can cause another type of resistance. As we see in Sections Effective treatment parameters differs for 2D and 3D cell arrangements and Optimal treatment parameters differs for 2D and 3D cell arrangements, there are values for the receptor’s kinetic parameters that prevent the cell from the regulatory effects produced by the binding of TNF. Therefore, when we consider heterogeneous populations by introducing variability in the kinetic parameters of the TNF receptor model, we observed that beyond a critical value of the parameter that controls variability most of the *effective treatments* that work in the homogeneous population fail to reduce tumor growth. Our hypothesis is that with high variability some of the cells could have kinetic parameters that make them insensitive to the treatment and thus they will produce the relapse, after the sensitive ones have been killed by the treatment. We found that this is the main cause of the non-optimal parameters sets found by the optimization techniques within heterogeneous populations.

Regarding the critical value for the control parameter, we found it is different for the 2D and the 3D cell arrangements, with a lower value in the latter case. We hypothesize that this difference may be due to two factors; first, because the space of *effective treatments* strategies shows to be more constrained in the 3D case. The second reason we propose is due to the differences in the total number of cells simulated in the 2D and 3D; while we set the same radius for the disc and the spheroid, the number of initial cells are 150 and 1000, respectively. Because variability is generated by sampling random parameters, a larger number of cells increases the probability of getting a set of kinetic parameters that make the cell insensitive to the TNF. We have shown that population variability can cause resistance. the higher the variability, the harder to find effective treatments. However, even in the cases of maximum variability analyzed the algorithms are able to find a few sets of candidate *effective treatments*.

## 1 Conclusions

Multi-scale modeling allows for gaining mechanistic insights in dynamic drug dosages and predict novel strategies for treatments. Even though, in the last few years, it has been great progress in the field [26], it is known that virtual drug screens seldom match with clinical trials results. Thus, we need to acknowledge that we are far from using these models at the patient’s bedside[16]. One of the improvements that would help close this gap would be to have simulations and optimizations that account for and embrace uncertainty. We have hereby presented a free to use, open-source framework that allows optimizing treatment strategies with varying levels of uncertainty. We tested the framework using a multi-scale model of cancer growth in different cell arrangements introducing population variability to show that population heterogeneity is critical, either be caused by the cells’ state, their parameters or the population size, and it affects the optimal parameter sets. We found that our model exploration workflow can find effective treatments in all the studied conditions, most importantly, our results show that cells’ spatial geometry, as well as, population variability should be considered when optimizing treatment strategies in order to find robust parameter sets.

## Conflict of Interest Statement

The authors declare that the research was conducted in the absence of any commercial or financial relationships that could be construed as a potential conflict of interest.

## Author Contributions

MPL conceived the work. MPL and AM implemented the model. MPL and CA implemented the workflow and the code for analyzing simulation results. MPL, Ca and AM prepared the manuscript. All authors edited, commented and agreed on the final version of the manuscript.

## Funding

This work has received funding from the EU Horizon 2020 RIA program INFORE under grant agreement No 825070 and the EU Horizon 2020 ICT program PerMedCoE under grant agreement No 951773.

## Acknowledgments

The authors acknowledge the technical expertise and assistance provided by the Spanish Supercomputing Network (Red Española de Supercomputación), as well as the computer resources used: the LaPalma Supercomputer, located at the Instituto de Astrofísica de Canarias and MareNostrum4, located at the Barcelona Supercomputing Center.

## Footnotes

1 https://github.com/bsc-life/spheroid-tnf-v2-emews.

## Notes

https://github.com/bsc-life/spheroid-tnf-v2-emews

### Competing Interest Statement

The authors have declared no competing interest.

